# Primary cilia regulate Meibomian glands development and dimensions without impairing lipid composition of the meibum

**DOI:** 10.1101/2022.05.16.492188

**Authors:** Céline Portal, Yvonne Lin, Varuni Rastogi, Cornelia Peterson, James Foster, Amber Wilkerson, Igor Butovich, Carlo Iomini

## Abstract

**Purpose:** Primary cilia regulate the development of various ectoderm-derived tissues, including the corneal epithelium, skin, hair follicle and sebaceous glands. We aimed to investigate their role in meibomian gland (MG) development.

**Methods:** Primary cilium ablation in MGs was achieved by crossing a floxed Ift88 mouse (*Ift88^fl/fl^*) with a mouse expressing the Cre recombinase under the keratin 14 (K14) promoter, to generate *K14-Cre;Ift88^fl/fl^* mice. MG morphology was evaluated by histology and immunostaining, as well as lipid staining and 2-photon microscopy on whole mount tarsal plates. MG lipid profiles were assessed by chromatography.

**Results:** We showed that most of MG cells are ciliated during early stages of MG development and that MG ciliated rate decreases throughout morphogenesis. In morphologically mature glands, only the MG central duct and ductules are ciliated, and meibocytes lose their cilia as they differentiate and become filled with lipids. Primary cilium ablation induces enlargement of MGs, dilation of the MG central duct, and an increased production of lipids, without dramatically changing the lipid profiles. In addition, primary cilia regulate MG elongation and the spatial distribution of proliferating and dying cells within MGs, without changing the total cell proliferation and death rates.

**Conclusions:** These findings indicate that primary cilia are not necessary for normal MG development. However, they promote MG enlargement and lipid production, suggesting that primary cilia could be an interesting target for treatments of ocular surface diseases involving MGs, like dry eye disease.

## INTRODUCTION

Meibomian glands (MGs) are holocrine glands located within the tarsal plates of the upper and lower eyelids. These modified sebaceous glands are composed of clusters of secretory acini, connected to a central duct by several shorter ductules. They release their secretory product, the meibum (composed of lipids, proteins and nucleic acids of the whole cell), at the eyelid margin, which is then spread onto the ocular surface, as the outermost layer of the tear film, with each blink (Bron et al., 2017; Knop et al., 2011). This lipid layer plays crucial protective roles for the ocular surface, as it functions as a lubricant for the eyelids during blinking, prevents tear overflow onto the lids, and reduces tear evaporation (Knop et al., 2011; McCulley & Shine, 2003).

Defective MGs lead to meibomian gland dysfunction (MGD), which has been defined as “a chronic, diffuse abnormality of the MGs, commonly characterized by terminal duct obstruction and/or qualitative/quantitative changes in the glandular secretion” (Nichols et al., 2011). Reduced lipid secretion may contribute to tear film instability and entry into the vicious cycle of dry eye disease (DED), which is among the most commonly encountered ophthalmic diseases and has a global prevalence ranging from 5 to 50% (Stapleton et al., 2017). DED is usually divided into two primary and non-mutually exclusive categories: aqueous deficient dry eye (ADDE) and evaporative dry eye (EDE) (Craig et al., 2017). MGD is considered the most common cause of EDE and the leading cause of DED (Craig et al., 2017; Foulks et al., 2012; Schaumberg et al., 2011). Indeed, EDE, alone or in combination with ADDE, affects more than 80% of patients with DED, while only 10% are attributed exclusively to ADDE (Kim et al., 2021). Despite recent development of therapeutic solutions targeting MGD (ocular lubricants, eyelid-warming devices, intense pulsed light) (Jones et al., 2017), these approaches primarily focus on relieving MG obstruction or replacing lipids, and there is an unmet need for treatments aimed at preventing MG atrophy and stimulating lipid production. However, this deficiency in current effective pharmacological targets is due to the very limited knowledge about molecular networks underlying MG development and renewal.

Human MGs form during embryonic development, between the third and the seventh month of gestation, corresponding to the sealed-lid phase of eyelid development (Knop et al., 2011). In mice, MG development begins at embryonic day 18.5 (E18.5) and continues postnatally (Nien et al., 2010). As in humans, MG development in mice occurs during the sealed-lid phase of eyelid development, which is indispensable for MG development (Meng et al., 2014; Wang et al., 2017).

Although it has been suggested that MG development shares similarities with the development of the pilosebaceous unit, which is composed of a hair follicle (HF) and its associated sebaceous glands (SG), the basic mechanisms underlying MG development and renewal remain poorly understood. Like HFs, MGs develop from the ectodermal sheet which invaginates into the mesoderm to form an anlage. Then, similar to the hair anlage of eyelashes, the meibomian anlage develops lateral outgrowths that later differentiate into ductules and sebaceous acini (Andersen et al., 1965). In murine development, the formation of an epithelial placode occurs at E18.5 which is followed by invagination in the mesenchyme and elongation of the placode, branching of the MGs beginning around postnatal day 5 (P5), and acquisition of their mature morphology by P15 (Nien et al., 2010).

Primary cilia are microtubule-based cellular organelles that originate from the basal body and extend from the plasma membrane. Intraflagellar transport (IFT), a bidirectional movement of protein particles along the axoneme, ensures the appropriate assembly and maintenance of cilia (Iomini et al., 2001; Ishikawa & Marshall, 2011; Kozminski et al., 1995; Rosenbaum & Witman, 2002; Taschner & Lorentzen, 2016). Dysfunction of the primary cilium produces a heterogeneous group of diseases called ciliopathies, some of which induce severe developmental defects, highlighting the crucial role of the primary cilium in tissue development (Reiter & Leroux, 2017). The primary cilium plays essential roles in the development of ectoderm-derived tissues including the skin, the corneal epithelium, and the pilosebaceous unit (Song & Zhou, 2020; Toriyama & Ishii, 2021). In particular, the primary cilium regulates corneal epithelial thickening through regulation of cell proliferation and vertical migration (Grisanti et al., 2016). In the skin, primary cilia limit hyperproliferation of keratinocytes in the epidermis (Croyle et al., 2011; Ezratty et al., 2011). Moreover, they are essential for HF morphogenesis, since primary cilium ablation leads to HF morphogenesis arrest (Chen et al., 2015; D Dai et al., 2013; Daisy Dai et al., 2011; Ezratty et al., 2011; Gao et al., 2008; Lehman et al., 2009; Liu et al., 2016), and abnormal primary cilia lead to dysregulation of the hair growth cycle (Lehman et al., 2008). Primary cilia also regulate SG development, and ablation of the primary cilium induces hyperplasia of sebaceous gland lobules (Croyle et al., 2011). Patients affected by Bardet-Biedl syndrome, an autosomal recessive ciliopathy, suffer from several cutaneous conditions including Keratosis pilaris and seborrheic dermatitis (Haws et al., 2019). Although the role of the primary cilium has been investigated in various ectoderm-derived tissues, its role in MG development, maintenance and function remains unknown.

In this study, we show that MG cells are ciliated in early stages of development, and meibocytes lose their primary cilium as they differentiate. We demonstrate that the primary cilium is required for regulating the central duct diameter and overall size of MGs. We propose a mechanism by which primary cilia determine the early MG cell patterning by controlling the spatial distribution of proliferating and dying cells within developing glands. These findings suggest cilia-mediated signaling pathways as possible therapeutic targets to counteract MGD.

## MATERIALS AND METHODS

### Mice

Mouse strains *Ift88^tm1Bky^* (here referred to as *Ift88^fl/fl^*) (Haycraft et al., 2007), B6N.Cg-Tg(KRT14-cre)1Amc/J (*K14-Cre*, Jackson Laboratory stock No 018964) (Dassule et al., 2000), and Gt(Rosa)26Sor(tm4(ACTB-tdTomato,-EGFP)Luo)/J (*mT/mG*, Jackson Laboratory stock No 007676) (Muzumdar et al., 2007) were maintained on mixed C57Bl/6, FVB and 129 genetic backgrounds. *Ift88* conditional knock-out (cKO) were generated by crossing *K14-Cre;Ift88^fl/+^* males with *Ift88^fl/fl^* females. Other allelic combinations than *K14-Cre;Ift88^fl/fl^* (cKO) were considered as controls (Ctrl). Mouse strain Tg(CAG-Arl13b/mCherry)1Kand Tg(CAG-EGFP/CETN2)3-4Jgg/KandJ (here referred to as Arl13b-mCherry;Centrin2-GFP) (Bangs et al., 2015) was purchased at Jackson Laboratory (stock No 027967). All animal procedures were performed in accordance with the guidelines and approval of the Animal Care and Use Committee at Johns Hopkins University and with the ARVO Statement for the Use of Animals on Ophthalmic and Vision Research.

### Histology and immunofluorescence staining

Upper and lower eyelids from P6, P8, P15 and P21 mice were dissected, fixed overnight in 4% paraformaldehyde (PFA) in PBS, and embedded in paraffin for histological analysis. Hematoxylin and eosin (HE) staining was performed following standard procedures. Sections were imaged with an Olympus slide scanner VS200 (Olympus, Center Valley, PA).

Eyelids from P3 mice were dissected, fixed for 2h in 4% PFA in PBS and embedded in optimal cutting temperature compound (OCT Tissue-Tek, Sakura Finetek, Torrance, CA). Cryosections were processed for ARL13B staining. After 10 min fixation with cold acetone (−20°C) and 20 min permeabilization with 0.2% Triton X-100 in PBS, sections were incubated with a rabbit anti-ARL13B antibody (17711-1-AP, ProteinTech Group, Rosemont, IL) in 2% BSA/0.1% Triton X-100/PBS overnight at 4°C. Sections were imaged with a Zeiss LSM880 confocal microscope (Zeiss, Iena, Germany).

### Fluorescent imaging of whole mount MGs

Eyelids from P1 and P8 *K14-Cre;Ift88^fl/fl^;mT/mG* mice (cKO) and *K14-Cre;Ift88^fl/+^;mT/mG* (Ctrl) littermates were dissected, fixed in 4% PFA in PBS for 1h and washed in PBS. During the fixation, most of the connective tissues and muscles covering the tarsal plate were manually removed. MGs were mounted in 90% glycerol and imaged with a LSM880 confocal microscope (for samples at P1) or a LSM710/NLO two-photon microscope (for samples at P8). Serial optical sections were acquired in 1 or 2 μm steps, through the entire MG. At P8, MG volume was quantified after 3D reconstruction of the z-stacks with Imaris (Bitplane, South Windsor, CT), and the number of acini per MG was manually counted.

### Oil Red O staining in whole mount MGs

Eyelids from P6, P8, and P21 mice were dissected, fixed in 4% PFA in PBS for 1h and washed in PBS. During the fixation, most of the connective tissues and muscles covering the tarsal plate were removed. MGs were stained for 1h in Oil Red O (ORO) solution (Electron Microscopy Sciences, Hatfield, PA) at room temperature (RT), rinsed with distilled H_2_O, mounted in 90% glycerol and imaged with an Olympus MVX10 dissecting scope (Olympus). MG size was determined by averaging the MG area of individual MGs measured with Fiji (Schindelin et al., 2012).

### Cilia localization

Eyelids from P3, P6, P8, P12 and P25 Arl13b-mCherry;Centrin2-GFP mice were dissected and fixed for 1h in 4% PFA/1% Triton X-100 (Mallinckrodt Pharmaceuticals, Staines-upon-Thames, United Kingdom) in PBS. Eyelids were then processed for K14 staining on whole MGs or embedded in OCT. For whole MGs, prior to staining, most of the connective tissues and muscles covering the tarsal plate were removed and tarsal plates were permeabilized for 1h with 2% BSA/1% Triton X-100/PBS at RT. MGs for whole mount samples and cryosections were stained with a rabbit anti-keratin 14 (K14) polyclonal antibody (1:1000, 905301, BioLegend, San Diego, CA). MG whole mounts and cryosections were imaged with a Zeiss LSM880 confocal microscope. Serial optical sections were acquired in 1 μm steps. After 3D reconstruction of the z-stacks with Imaris, MGs were outlined using the Surfaces tool, and primary cilia and basal bodies were counted in MGs using the Spots tool. Centrin2-GFP labeled-centrioles were considered as a single basal body when less than 2 μm apart. The percentage of basal bodies associated with a primary cilium in MGs was determined by normalizing the number of primary cilia to the number of basal bodies.

### In vivo cell proliferation assay

Mice at P6 and P21 received a single intraperitoneal injection of 50 mg/kg EdU (EdU-Click 594, baseclick, Germany) and were sacrificed after 6h, as described elsewhere (Sun et al., 2018). Eyelids were dissected and snap frozen in OCT. Twenty micron cryosections were processed following manufacturer instructions (EdU-Click 594, baseclick, Germany). Briefly, sections were fixed for 15 min with 4% PFA in PBS, permeabilized for 20 min with 0.5% Triton X-100 in PBS and then stained for 30 min with the reaction cocktail in the dark at RT. MGs were localized using the mT/mG reporter or stained using a rabbit anti-K14 polyclonal antibody (1:500, 905301, BioLegend). Nuclei were counterstained with DAPI. Sections were imaged with a Leica DMI6000 microscope equipped with a Yokogawa confocal spinning disc or a Zeiss LSM880 confocal microscope. For each cryosection, serial optical sections were acquired in 2.45 μm steps. For each mouse, serial cryosections were processed to acquire the entirety of the MGs. After 3D reconstruction of the z-stacks with Imaris, MGs were outlined using the Surfaces tool, and EdU-positive nuclei and DAPI-positive nuclei were counted in each MG using the Spots tool. Quantification was performed on the MGs located in the center of the upper eyelid. The cell proliferation rate per MG was determined by normalizing the number of EdU-positive nuclei to the number of DAPI-positive nuclei. Quantification of the EdU-positive and DAPI-positive nuclei was also performed, separating the proximal part (from the eyelid margin to the middle of the gland) and the distal part (from the middle of the gland to the tip) of the MGs.

### Cell death TUNEL assay

Eyelids from P21 mice were dissected, fixed overnight with 4% PFA in PBS and embedded in paraffin. Sections were processed for apoptosis immunofluorescent staining using the *In situ* Cell Death Detection Kit, TMR Red (Roche Applied Science, Mannheim, Germany) as previously described (Ibrahim et al., 2014). Sections were permeabilized for 15 min with 10 μg/mL proteinase K in 10 mmol/L Tris/HCl (pH 7.4) at RT, and then stained for 1h with the reaction mixture at 37°C in the dark. Nuclei were counterstained with DAPI. Sections were imaged with a Zeiss LSM880 confocal microscope. TUNEL-positive and DAPI-positive nuclei were counted on 3 serial sections, 20 μm apart from each other. The cell death rate per MG was determined by normalizing the number of TUNEL-positive nuclei to the number of DAPI-positive nuclei.

### Lipid analysis

Meibomian lipids were extracted from surgically excised mouse tarsal plates (4 from each mouse) at 4°C using three sequential extractions with a chloroform:methanol (3:1, vol:vol) solvent mixture. The extracts (3 x 1mL) were pooled, and the solvent was evaporated under a stream of compressed nitrogen at 37°C. The oily residue was redissolved in 1 mL of LC/MS quality iso-propanol and stored in a nitrogen-flushed, crimper-sealed HPLC 2-mL autoinjector vial at −80°C before the analyses.

The gradient and isocratic reverse-phase liquid chromatography/high resolution time of flight atmospheric pressure chemical ionization mass spectrometry (LC/MS) analyses were conducted using, correspondingly, an Acquity UPLC C18 (1 mm x 100 mm; 1.7 μm particle size) and an Acquity UPLC C8 (2 mm x 100 mm; 1.7 μm particle size) columns (both from Waters Corp., Milford, MA, USA) as described in detail in our earlier publications for mouse and human meibum (I A Butovich et al., 2019, 2021; Igor A. Butovich & Suzuki, 2021). Between 0.5 and 1.0 μL of the sample solution was injected per experiment. A Waters Acquity M-Class binary UPLC system (Waters Corp.) was operated at a 20 μL/min flow rate. The analytes were eluted using acetonitrile/iso-propanol isocratic solvent mixtures with 5% of 10 mM ammonium formate as additive. The analytes were detected using a high-resolution Synapt G2-Si QToF mass spectrometer (equipped with a ZSpray interface, an IonSabre-II atmospheric pressure chemical ionization (APCI) ion source and a LockSpray unit (all from Waters Corp.)). All the experiments were conducted in the positive ion mode. Most of the lipids were detected as (M + H)^+^ and (M + H – H_2_O)^+^ adducts. However, some of the, for example triacylglycerols and cholesteryl esters, underwent spontaneous in-source fragmentation producing (M + H - fatty acid)^+^ species. The major lipid analytes were identified using the EleComp routine of the MassLynx v.4.1 software package (Waters Corp.), though detailed analyses of the mouse MG lipidomes have not been performed at this time.

The total lipid production by MGs was estimated on the basis of the total ion chromatograms of their lipid extracts recorded in isocratic LC/MS APCI experiments as described recently (Igor A. Butovich & Suzuki, 2021). The unbiased, untargeted analysis of the lipidomic data was conducted using Progenesis QI and EZinfo software packages (Waters Corp.). The Principal Component Analysis (PCA-X) approach was used to evaluate the differences between the control and cKO samples.

### Statistics

Data were presented as mean±SD. Mann Whitney test, Wilcoxon signed rank test, and one-way ANOVA with post-hoc Tukey HSD test were performed with the online web statistical calculators https://astatsa.com/ or RStudio (Team, 2021). A *P* value < 0.05 was considered significant.

## RESULTS

### Primary cilium ablation leads to larger MGs

To determine if primary cilia are involved in MG development, we specifically ablated *Ift88*, a subunit of the intraflagellar transport machinery required for cilia assembly and maintenance (Pazour et al., 2000) in MGs, by generating the *K14-Cre;Ift88^fl/fl^* conditional knock-out (here referred as cKO). In this mouse, the *Ift88* gene is excised in all epithelial cells expressing K14, including the MGs. The *K14-Cre* recombinase expression was followed by using the mT/mG reporter mouse line (Muzumdar et al., 2007) (**Supplement figure 1**). In the *K14-Cre;Ift88^floxed^;mT/mG* transgenic line, the Cre-dependent excision of a cassette expressing the red-fluorescent membrane-targeted tdTomato (mT) drove the expression of a membrane-targeted green fluorescent protein (mG) in K14-expressing tissues, including MGs (**Supplement figure 1**). To monitor primary cilium ablation in MGs, we labeled primary cilia using an anti-ARL 13B antibody. In control mice, virtually all the cells in MGs were ciliated at P3 while in cKO mice, cilia were absent in most of the MG cells (**Supplement figure 1**).

As previously published, no evident defects were visible in newborn cKO mice and eyelid opening occurs around P13 in both control and cKO mice (Croyle et al., 2011; Grisanti et al., 2016). Any potential defects in MG development observed in the cKO mutant would not be due to defect in eyelid fusion/opening, since eyelid fusion is a mandatory step in MG morphogenesis (Wang et al., 2017). However, we noticed the presence of multifocal, white, granular to chalky, seborrheic debris along the eyelid margins of most of adult cKO mice, which was never observed in control mice (**Figure 1A**). MG were stained with ORO to assess their morphology during development. There was no significant difference observed between males and females of the same genotype (**Supplement figure 2**), and both sexes were pooled for the entire study. MGs were visible in both the upper and the lower eyelids of both control and cKO mice (**Figure 1B**), and the number of glands was the same in both genotypes (**Figure 1D**) at P6 and P8. However, MGs were 29% and 21% larger in cKO mice compared to control mice at P6 and P8, respectively (**Figure 1D**). At P21, when the MGs had reached their mature morphology, the margins between individual MG was indistinct, and obtaining the area of individual glands was not possible (**Figure 1C**). Overall, MGs in the cKO tarsal plates appeared more densely arranged than in control eyelids. When evaluated by ORO, there were negative staining foci between the MGs and the HFs in the controls, whereas the tarsal plate of cKO mice were characterized by diffuse ORO-positive acini. At the cellular level, meibocytes seemed similar between cKO and control mice at P6 and P21 (**Figure 1E**). Collectively, primary cilium ablation did not suggest impaired MG maturation but promoted the development of larger MGs.

**Figure 1:**
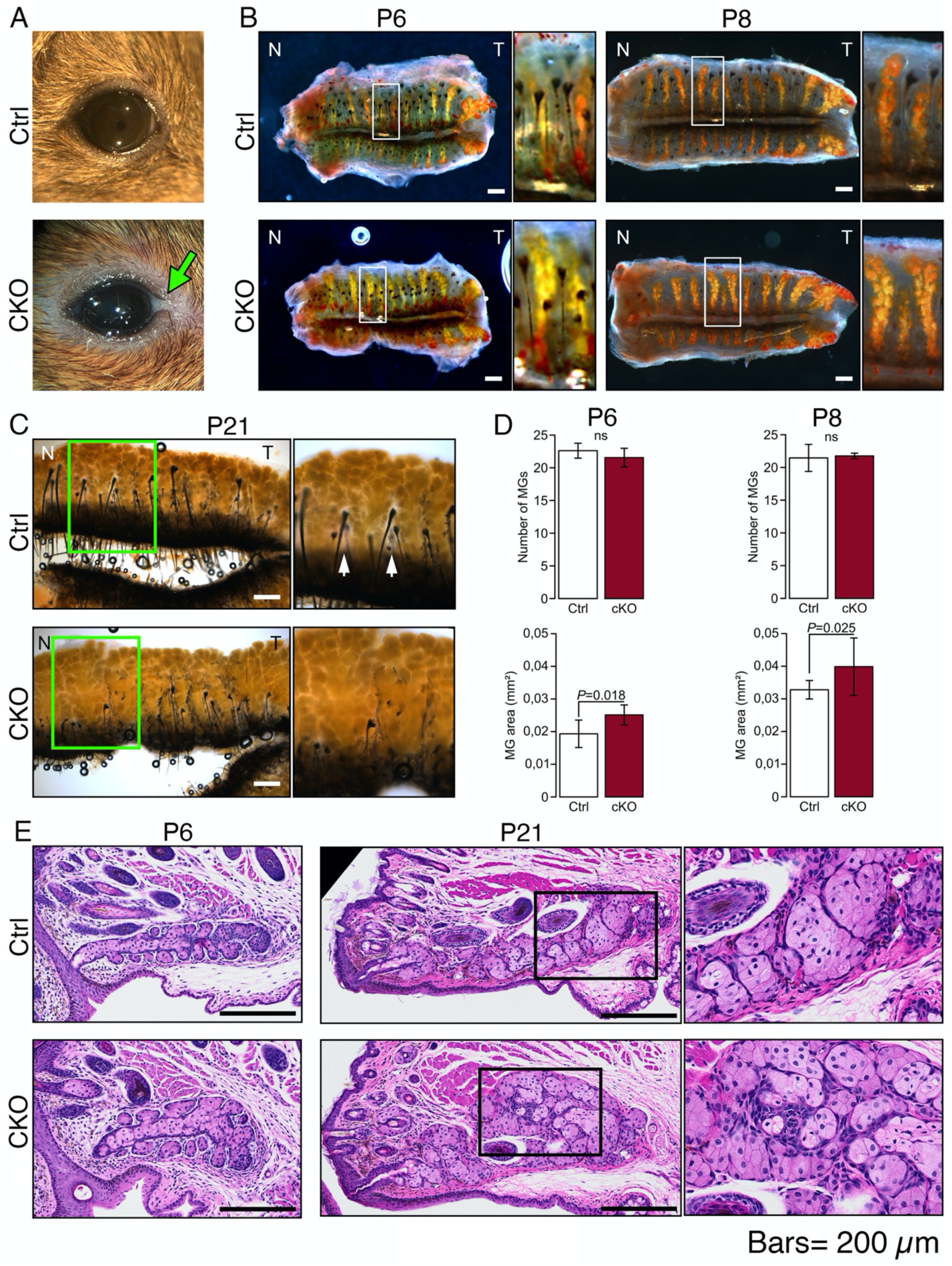
Primary cilium ablation leads to larger MGs. (**A**) Representative pictures of control and cKO adult (6 months) eyes. Arrow indicates white deposit, only observed in cKO mice. (**B-C**) Representative images of tarsal plates stained with ORO staining at P6, P8, and P21. Boxed regions indicate the areas shown at higher magnification. Scale bar; 200 μm; N, nasal; T, temporal. (**D**) Number of MGs and MG size were quantified at P6 and P8 (n=20 controls and 4 cKO mice at P6; n=13 controls and 9 cKO mice at P8). Per mouse, MG area was determined by averaging the MG area of all individual MGs in the upper and lower eyelids. Data were presented as mean±SD. Statistical significance was assessed using Mann Whitney test. ns, non-significant, *P*≥0.05. (**E**) Representative images of control and cKO MGs stained with HE at P6 and P21. Boxed regions indicate the areas shown at higher magnification. Scale bar; 200 μm.

### Primary cilium ablation induces an increased production of lipids

To further characterize the effect of primary cilium ablation on meibocyte maturation, we performed lipidomic analysis in adult MGs. Lipid profiles were assessed by high-resolution mass spectrometry (MS), MS/MS and isocratic and gradient reverse-phase ultra-high performance liquid chromatography on lipids extracted from tarsal plates. Approximately 150 analytes with unique combinations of retention times (RTs) and mass-to-charge (*m/z*) ratios were detected, and representative elution profiles from a control mouse are presented in **Figures 2A-C**. The PCA-X analysis produced no obvious clustering of either the control or cKO samples (**Figure 2D**), implying that their chemical compositions were similar to each other. However, analysis of samples for a set of 15 major lipid species from the wax ester (WE) and the cholesteryl ester (CE) families showed a ~2-fold increase in the amount of produced lipids in the cKO mutant (**Figure 2E**). Mice heterozygous for the mutation demonstrated a subtler effect than their homozygous littermates.

**Figure 2:**
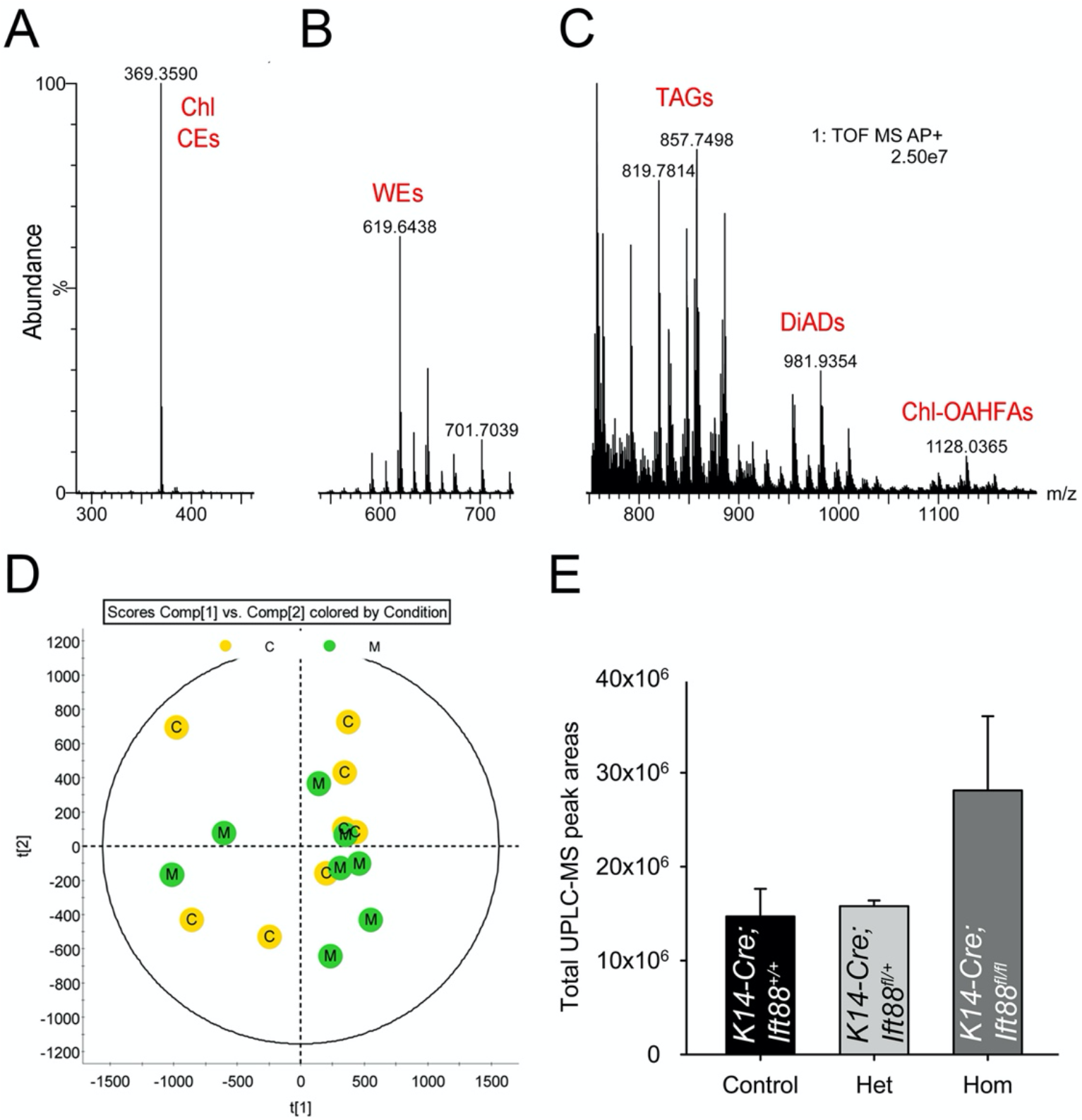
Reverse phase liquid chromatography/high resolution time of flight atmospheric pressure chemical ionization mass spectrometry (LC/MS) analysis of mouse meibomian lipids conducted in the positive ion mode revealed close similarity in the lipidomes of control and cKO mice, but a twofold increase in the total lipid content of the latter. (**A**) An analytical signal of free cholesterol (Chl) and cholesteryl esters (CEs) from a control mouse. (**B**) An observation spectrum of the pool of meibomian wax esters (WEs) from a control mouse. (**C**) An observation spectrum of the pools of triacylglycerols (TAGs), α,ω-diacylated diols (DiADs), and cholesteryl esters of (*O*)-acylated ω-hydroxy fatty acids (Chl-OAHFAs) from a control mouse. (**D**) A scores plot generated using Principal Component Analysis (PCA) for the control (C, yellow dots) and cKO (M, green dots) LC/MS data demonstrated a strong overlap of the control and mutant samples of meibomian lipids with no clear clustering of the samples of different types, indicating their close biochemical compositions. (**E**) However, the primary cilium ablation led to a higher overall lipid production in the tarsal plates of cKO mice compared with the lipid content of control mice (n=8 for Ctrl, n=3 for Het, n=8 for cKO). In this figure, Hom stands for homozygotes and corresponds to the cKO mice (*K14-Cre;Ift88^fl/fl^*), Het corresponds to the heterozygote mice (*K14-Cre;Ift88^fl/+^*) and control corresponds to all the other genetic combinations possible.

### MGs are ciliated throughout their development and lose their cilia as meibocytes mature

Primary cilia are indispensable for adipocyte differentiation (Zhu et al., 2009); however, we showed that the loss of primary cilia does not dramatically affect meibocyte differentiation. In adipocytes, a variation of cilia size during early stages of adipocyte differentiation and the loss of primary cilia during adipocyte maturation has been reported (Forcioli-Conti et al., 2015, 2016; Marion et al., 2009). To our knowledge, primary cilia have not been described in MGs. To determine the spatiotemporal distribution of primary cilia during MG development, we used the Arl13b-mCherry;Centrin2-GFP mouse (Bangs et al., 2015) in which ARL13B, a widely accepted ciliary marker (Caspary et al., 2007), is fused with mCherry labeling primary cilia in red, and Centrin2, one of the classical basal body associated centrins (Ruiz et al., 2005), is fused with GFP labeling basal bodies in green.

On whole mount tarsal plates, we noticed considerable variability in MG size, especially MGs on the temporal aspect of the eyelid which were larger with a less linear morphology than central or nasal MGs (**Figure 1B**). To overcome this bias, we decided to number MGs starting from the temporal side and to focus the rest of the study on MGs localized in the center of the eyelid (MG #4 through #8) (**Figure 3A**). We first determined the spatiotemporal distribution of primary cilia during MG development. At P3, primary cilia were diffusely visible along the MG anlage, essentially in the center of the gland (**Figure 3C-D**). In particular, primary cilia were located along the apical aspect of the basal cells of the MG and oriented toward the center of the gland (**Figure 3D**). At P3, more than 70% of the cells were ciliated (**Figure 3G**). A similar proportion of MG cells were ciliated during early MG branching at P6 and P8 (**Figure 3G**). At P6, primary cilia were visible in the elongating distal tip, the developing duct, and the budding acinus (**Figure 3E**). As MGs continued to mature, the percentage of basal bodies associated with a primary cilium significantly decreased at P12 and P25 (**Figure 3G**). The percentage of ciliated cells was similar in the duct and acini of MGs at P12 (**Figure 3H**), but, interestingly, primary cilia were no longer present in P25 acini (**Figure 3F, 3H**). At P25, primary cilia were only visible in the central duct and connecting ductules of the morphologically mature MGs (**Figure 3F**). Thus, meibocytes lose their primary cilium as they mature and become filled with lipids.

**Figure 3:**
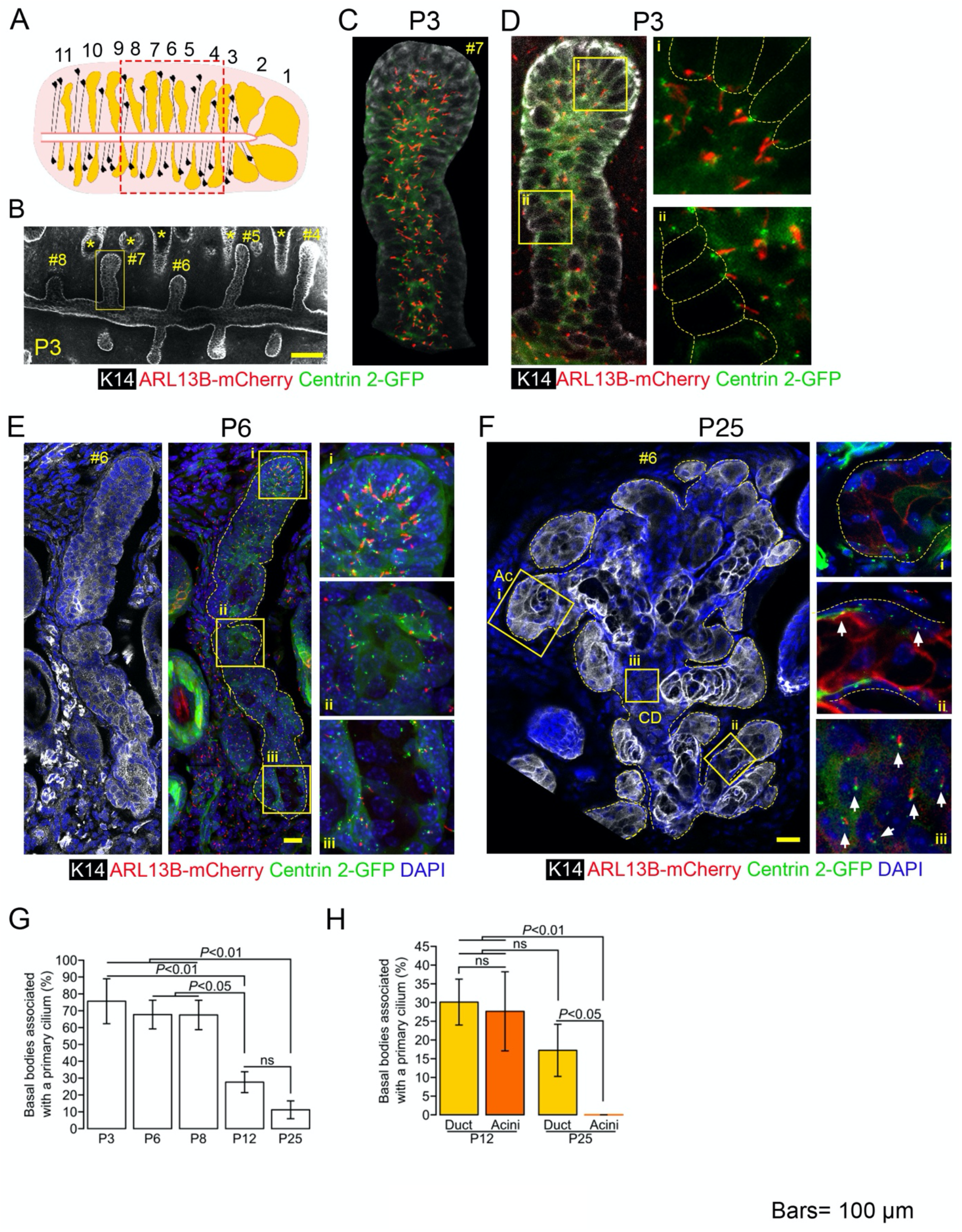
Most MG cells were ciliated in early stages of MG development, but meibocytes lost their cilium as they matured. (**A**) To facilitate comparison of similar MGs, MGs were numbered from the temporal side (MG#1) to the nasal side (MG#11). The boxed area indicates the region in which MGs were studied for all the following experiments (**B**) Whole mount tarsal plate at P3 imaged by confocal microscopy. MGs were stained with an antibody against K14 (in white). HFs (marked with *) were also stained but were easily distinguishable from MGs due to the presence of a hair shaft. The boxed region indicates the MG shown at higher magnification in **C** and **D**. Scale bar; 100 μm. (**C**) 3D reconstruction with Imaris of MG#7 In Arl13b-mCherry;Centrin2-GFP mice, mCherry labels primary cilia in red, and GFP labels basal bodies in green. At P3, cilia were visible all along the MG. Scale bar; 100 μm. (**D**) Optical section picked in the center of MG#7. Basal bodies (in green) and primary cilia (in red) were localized on the apical side of MG basal cells (outlined with a yellow dotted line). Scale bar; 100 μm. (**E**) Representative MG longitudinal section at P6. MGs (outlined by a yellow dotted line) were stained with an antibody against K14 (in white), nuclei were stained with DAPI (in blue), basal bodies were labeled with GFP (in green) and primary cilia were labeled with mCherry (in red). MG cells are ciliated all along the MG, including in the distal tip of the gland (i), the forming acini (ii) and the forming central duct (iii). Scale bar; 100 μm. (**F**) Representative longitudinal section of a morphologically mature MG at P25. MGs (outlined by a yellow dotted line) were stained with an antibody against K14 (in white), nuclei were stained with DAPI (in blue), basal bodies were labeled with GFP (in green) and primary cilia were labeled with mCherry (in red). No primary cilia were visible in the acini (i) but primary cilia (arrows) were still present in ductules (ii) and the central duct (iii). Scale bar; 100 μm; Ac, acini; CD, central duct. (**G**) Quantification of the percentage of basal bodies associated with a primary cilium throughout development (n=3 for each age). Data were presented as mean±SD. Statistical significance was assessed using ANOVA one-way with post-hoc Tukey HSD test. ns, non-significant, *P*≥0.05. (**H**) Quantification of the percentage of basal bodies associated with a primary cilium in the acini and in the duct of MGs at P12 and P25 (n=3 for each age). Data were presented as mean±SD. Statistical significance was assessed using Mann Whitney test (Ctrl vs cKO) and Wilcoxon signed rank test (acini vs duct). ns, nonsignificant, *P*≥0.05.

### Primary cilium ablation induces a mislocalization of proliferating and dying cells within MGs

Several studies suggest that primary cilium resorption is necessary for cell proliferation, and have demonstrated either reduction or loss of primary cilia in various epithelial neoplasms (reviewed in (Goto et al., 2017; Halder et al., 2020; Sánchez & Dynlacht, 2016)). We therefore hypothesized that the larger MGs induced by primary cilium ablation could be due to an increased meibocyte proliferation. Cell proliferation rates were assessed by counting the number of EdU-positive cells 6h after EdU injection and normalized to the total number of cells. EdU-positive and DAPI-positive cells were counted on serial sections to obtain proliferation rates in the full MGs. Cell proliferation rates were assessed at P4, P6, and P21 when MGs are elongating, starting to branch, and reaching their mature morphology (Nien et al., 2010), respectively. Cell proliferation rates in MGs were not significantly different between cKO and control mice, at any time point assessed (**Figure 4D**). However, we observed an accumulation of proliferating cells in the distal tip of control MGs at P4 and P6 that was not present in cKO samples (**Figure 4A-B**). Thus, we quantified the cell proliferation rate specifically in the distal half of MGs, from the center of the gland to the distal tip, and in the proximal half of MGs, from the eyelid margin to the center of the gland. At P4 and P6, there were significantly more proliferating cells in the distal half than in the proximal half of control MGs, whereas cell proliferation rates between the distal and the proximal halves did not differ in the cKO MGs (**Figure 4E**). At P21, there were significantly more proliferating cells in the distal half of control MGs compared to the proximal aspect, despite the distal enrichment of proliferating cells in the MGs of control mice not being a feature appreciated on EdU-stained sections (**Figures 4C and E**). Similar to the earlier time points, there was no significant difference between cell proliferation rates in the distal and proximal aspects of cKO MGs (**Figure 4E**). Thus, primary cilium ablation does not regulate the cell proliferation rate during MG development, but instead defines the localization of proliferating cells within MGs.

**Figure 4:**
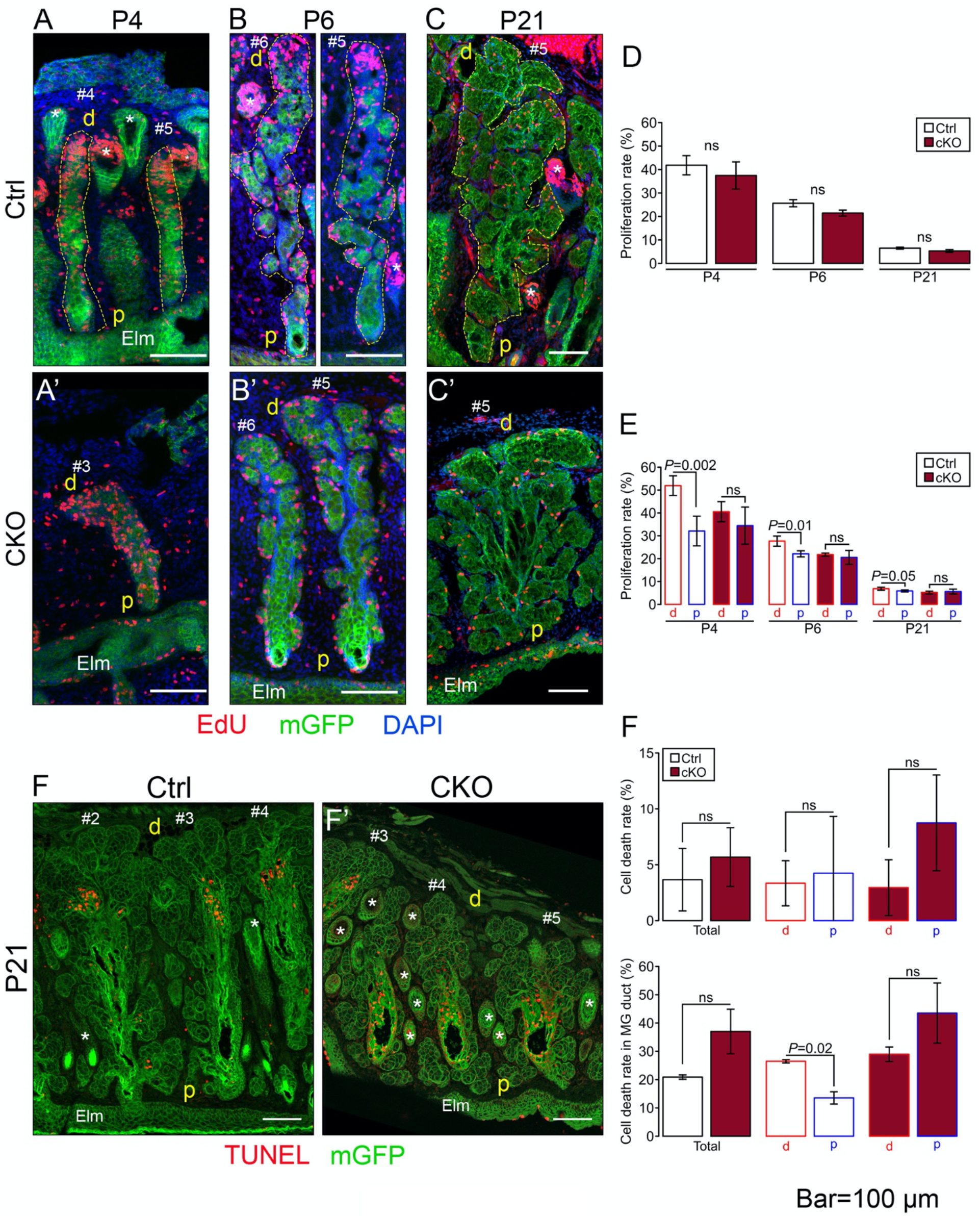
Primary cilium ablation did not change the overall rates of proliferation and dying cells in MGs but led to an abnormal distal/proximal localization of the proliferating and dying cells within MGs. (**A-C**) Cell proliferation was assessed by EdU staining in MGs at P4, P6 and P21 in control (**A**, **B**, and **C**) and cKO (**A’**, **B’**, and **C’**) mice, respectively. Scale bar; 100 μm; *, hair follicle; Elm, eyelid margin; d, distal; p, proximal. (**D-E**) Proliferation rates were quantified at P4, P6 and P21 in the full MGs (**D**), and specifically in the proximal (p) half (from the eyelid margin to the center of the gland) and the distal (d) half (from the middle to the tip of the gland) of MGs (**E**). Proliferation rates were determined by normalizing the number of EdU-positive nuclei to the total number of nuclei stained by DAPI (n=4/group). Data were presented as mean±SD. Statistical significance was assessed using Mann Whitney test (Ctrl vs cKO) and Wilcoxon signed rank test (distal vs proximal). ns, non-significant, *P*≥0.05. (**F**) Cell death was assessed by TUNEL staining in MGs at P21 in control (F) and cKO (F**’**) mice, respectively. Scale bar; 100 μm; *, hair follicle; Elm, eyelid margin; d, distal; p, proximal. (**G**) Cell death rates were quantified at P21 in the full MGs (*top graph*) and specifically in the central duct (*bottom graph*). Cell death rates were determined by normalizing the number of TUNEL-positive nuclei to the total number of nuclei stained by DAPI (n=3/group). Data were presented as mean±SD. Statistical significance was assessed using Mann Whitney test (total, Ctrl vs cKO) and Wilcoxon signed rank test (distal vs proximal). ns, non-significant, *P*≥0.05.

Since we identified areas of enrichment of proliferating cells during MG development, especially in the earlier time points, we next explored whether this correlated to primary cilium localization within MGs. To determine if there were areas richer in primary cilia within MGs, we used the Arl13b-mCherry;Centrin2-GFP mouse to locate primary cilia. The number of basal bodies associated with a primary cilium was not significantly different between the distal and proximal halves of MGs at P3, P6, and P8 (**Supplement figure 3**).

Since there was no significant difference in cell proliferation rates between control and cKO mice, we speculated whether the larger MGs of the cKO mice were due to a decreased cell death rate. We quantified the percentage of dying cells in morphologically mature MGs at P21 by TUNEL staining. First, we counted dying cells in all aspects of the MG, including the central duct, the ductules and the acini. The cell death rate was not significantly different between cKO and control mice, and the cell death rate in the distal half was not significantly different from the proximal half in either the control or the cKO mice (**Figure 4G *top***). However, it appeared that the dying cells were predominantly located in the central duct of MGs (**Figure 4F**), so we also quantified the cell death rate specifically in the central duct. Despite there being no significant differences in total cell death rates between control and cKO MGs, specifically within the central duct, the cell death rate was significantly higher in the distal half compared to the proximal half in control but not in cKO MGs (**Figure 4G bottom**). Thus, similar to the conclusions drawn from our proliferation assays, ablation of the primary cilium does not regulate the cell death rate, but rather defines the localization of dying cells within MGs.

### Primary cilium regulates MG elongation and MG central duct width but not MG branching

Since primary cilia regulate the spatial distribution of proliferating and dying cells within MGs, we next aimed to assess how the primary cilium affects the overall cell distribution and 3D morphogenesis of MGs. Thus, we took advantage of the mT/mG fluorescent reporter to assess these aspects at the cellular resolution. At P1, meibomian anlages looked similar between cKO and control mice (**Figure 5A-B**). However, the cells of the cKO MGs appeared more randomly distributed when compared to the cells in the center of the anlage in the control, which appeared uniformly aligned. By P3, MGs appeared more elongated in the control than in cKO mice (**Figure 5C-E**). MG width was significantly increased in the cKO compared to the control (**Figure 5F**), and MG length was significantly reduced in the cKO compared to the control at P3 (**Figure 5G**). Moreover, the ratio of the number of basal cells compared to the number of cells in the center of the gland was significantly reduced in cKO mice, indicating that there were more cells in the central part of cKO MGs (**Figure 5H-I**). As MGs continued to develop, we noticed an enlargement of the central duct in cKO MG, which was significantly increased at P6 and P21 (**Figure 6A-B**). At P8, MGs were about 2 times larger in volume in cKO than in control mice (**Figure 6C, D**). However, the number of acini was not significantly different between both genotypes (**Figure 6D**). Thus, primary cilia promote the segregation of proliferating cells at the distal tip of the growing MGs, which in turn ensures a balanced growth in length and width of the MG duct.

**Figure 5:**
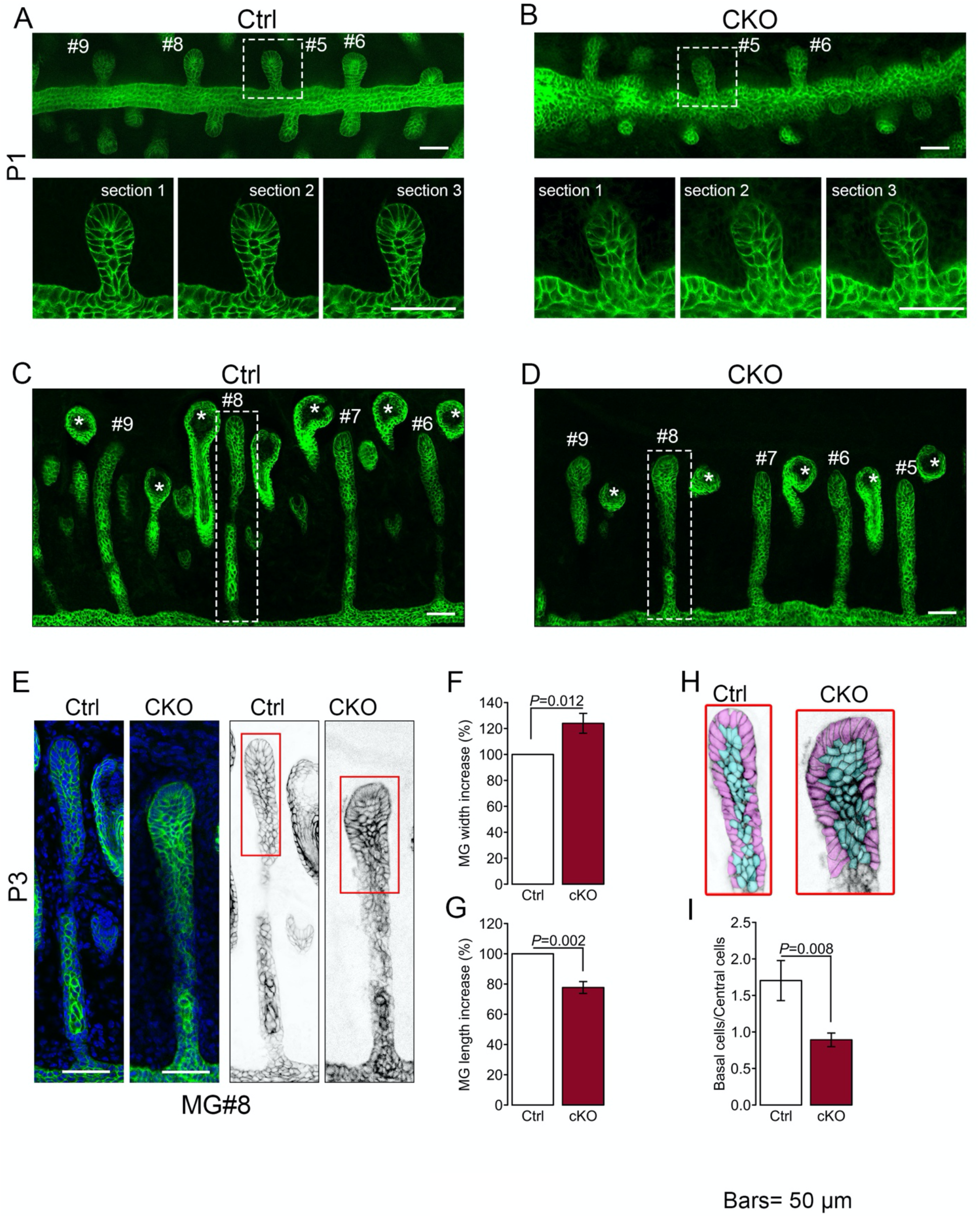
Primary cilium regulates MG elongation. (**A-B**) Representative overview of whole mount tarsal plates at P1 imaged by confocal microscopy. MGs were visualized by the mG fluorescent reporter. Boxed regions indicate MGs shown at higher magnification, for which serial optical sections are displayed. Scale bar; 50 μm. (**C-D**) Representative overview of whole mount tarsal plates at P3 imaged by confocal microscopy. MGs were visualized by the mG fluorescent reporter. Boxed regions indicate MGs shown at higher magnification in **E**. Scale bar; 50 μm. MG width (**F**) and MG length (**G**) of cKO mice normalized to control (n=3 mice/group). Data were presented as mean±SD. Statistical significance was assessed using Mann Whitney test. (**H**) Basal cells (in purple) and central cells (in blue) were manually color coded and then counted. (**I**) The ratio between basal cells and central cells was calculated (n=3 mice/group). Data were presented as mean±SD. Statistical significance was assessed using Mann Whitney test.

**Figure 6:**
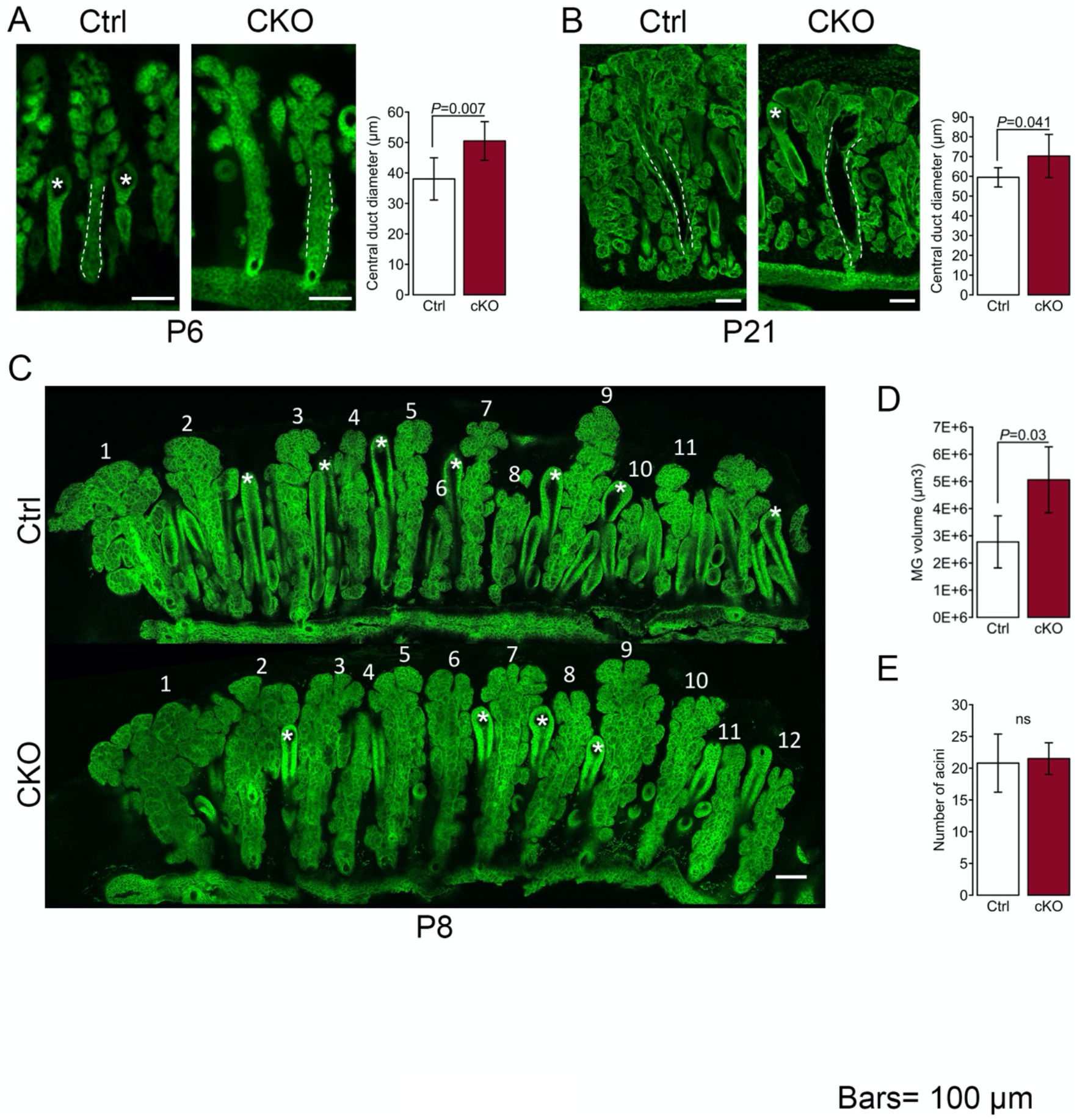
Primary cilium ablation induces dilation of the MG central duct but does not affect MG branching. (**A-B**) Representative MG longitudinal sections at P6 (**A**) and P21 (**B**). MGs were visualized by the mG fluorescent reporter, and MG central duct diameter (outlined by white dotted lines) was measured (n=7 controls and 7 cKO at P6; n=7 controls and 8 cKO at P21). Data were presented as mean±SD. Statistical significance was assessed using Mann Whitney test. Scale bar; 100 μm. (**C**) Representative overview of whole mount tarsal plates at P8 imaged by 2-photon microscopy. Scale bar; 100 μm. (**D**) MG volume at P8 was quantified after 3D reconstruction of z-stacks using Imaris (n=5 controls and 4 cKO). (**E**) MG branching was assessed by counting the number of acini per gland (n=5 controls and 4 cKO). Data were presented as mean±SD. Statistical significance was assessed using Mann Whitney test. ns, nonsignificant, *P*≥0.05.

## DISCUSSION

In this study, we have shown that meibocytes lose their primary cilium as they mature. This finding is similar to adipocytes, which are ciliated during early stages of adipogenesis, and then lose their cilia as they mature and their cytoplasm is expanded by lipids (Forcioli-Conti et al., 2015; Marion et al., 2009). However, primary cilium ablation in meibocytes leads to the opposite effects to those observed following primary cilium ablation in adipocytes, highlighting that the roles of primary cilia are cell and tissue-specific. Whereas ablation of the primary cilium in meibocytes does not seem to impair differentiation, the absence of the primary cilium in preadipocytes prevents differentiation into adipocytes (Zhu et al., 2009). Here, we have demonstrated the presence of ciliated cells in the morphologically mature MG ductules and a paucity of ciliated cells within the acini. This critical finding correspond to previous studies which have established that stem cells are ciliated (Shimada & Kato, 2022) and, further, that the stem cell niche is localized to the ductules, at the point in which the central duct transitions to the acini, in adult MGs (Parfitt et al., 2016). As primary cilia regulate various signaling cascades during development, it would be of particular interest to investigate which pathways are specifically affected during MG morphogenesis.

The tissue-specific roles of primary cilia also include regulation of gland branching. Surprisingly, differences in MG branching were not observed in the current study, whereas abnormal branching has been described in various glands in which primary cilia have been ablated. Notably, the ablation of primary cilia has been shown to induce formation of excess SG lobules (Croyle et al., 2011). Primary cilium ablation of the mammary gland, however, reduces branching and promotes the formation of smaller glands (McDermott et al., 2010).

Since we have shown that primary cilium ablation results in larger MGs with increased lipid production but similar lipid profiles when compared to control mice, we hypothesize that primary cilia may be a novel therapeutic target for ocular surface diseases which result from deficiencies in lipid synthesis or secretion such as MGD and DED. The current clinical interventions for DED and MGD are primarily replacement strategies, using artificial tears to maintain ocular surface hydration and lubrication (Jones et al., 2017). However, the efficacy of this approach in alleviating dryness symptoms is transient, and patients must apply these topical preparations several times a day. Expanding treatment strategies to address alternative causes of symptoms by directly targeting lipid production, has the potential to not only relieve ocular surface discomfort, but also improve the quality of life in affected patients.

In conclusion, we have shown that while primary cilia are not essential for MG development, their demonstrated capacity to increase lipid production and to regulate the spatial distribution of proliferating and dying cells within the MG may present a target for the development of new drugs for MGD and DED treatment. Further investigations are needed to identify specific signaling pathways trafficking through the cilium and whether primary cilium ablation in models of MGD could reverse MG atrophy and restore lipid production.

## ACKOWLEDGEMENTS

The authors thank Q Liu for his involvement during the early stages of this project, Hoku West-Foyle from the Microscope Facility at Johns Hopkins for his technical assistance, the reference histology core facility at Johns Hopkins University for the paraffin sections, and the members of the Wilmer Cornea Group for their helpful input and discussions. This work was supported by grants from the National Eye Institute, National Institute of Health (EY030661 to CI, EY024324 and EY027349 to IB); by a core grant to the Wilmer Eye Institute (EY001765), by a grant from the Office of Director, NIH (S10RR024550 to S.C. Kuo, Microscopy Facility at Johns Hopkins University); by a grant from the Eisinger Family; by a Wilmer Eye Institute Seed Fund to CI; and by an unrestricted grant from RPB to the Wilmer Eye Institute.

---

**Supplement figure 1: Genetic deletion of Ift88 in K14-expressing cells leads to primary cilium ablation in MGs.** (**A**) Breeding strategy to generate conditional ciliary mutant and visualize Cre expression. (**B**) Representative MG sections of control and cKO mice at P3. MGs (surrounded by white dotted line) express the mG reporter. Primary cilia were stained with an anti-ARL13B antibody, respectively. Scale bar; 10 μm.

**Supplement figure 2: MG size is not sex-dependent during MG development.** MGs were stained with ORO in whole mount tarsal plates females in control (**A**) and cKO (**B**) mice at P8. Per mouse, MG area was determined by averaging the MG area of all individual MGs in the upper and lower eyelids. Data were presented as mean±SD (n=6 females and 7 males for control mice; n=4 females and 5 males for cKO mice). Statistical significance was assessed using Mann Whitney test. ns, non-significant, *P*≥0.05.

**Supplement figure 3: Primary cilia are homogeneously distributed with MGs during early steps of MG development.** The percentage of basal bodies associated with a primary cilium was quantified in the distal half and in the proximal half of MGs at P3, P6 and P8 (n=3 for each age). Data were presented as mean±SD. Statistical significance was assessed using Wilcoxon signed rank test (distal vs proximal). ns, non-significant, *P*≥0.05.

## Notes

### Competing Interest Statement

The authors have declared no competing interest.

