## Supplementary figures and images for "Primary cilia regulate Meibomian glands development and dimensions without impairing lipid composition of the meibum"

### Supplemental Figures

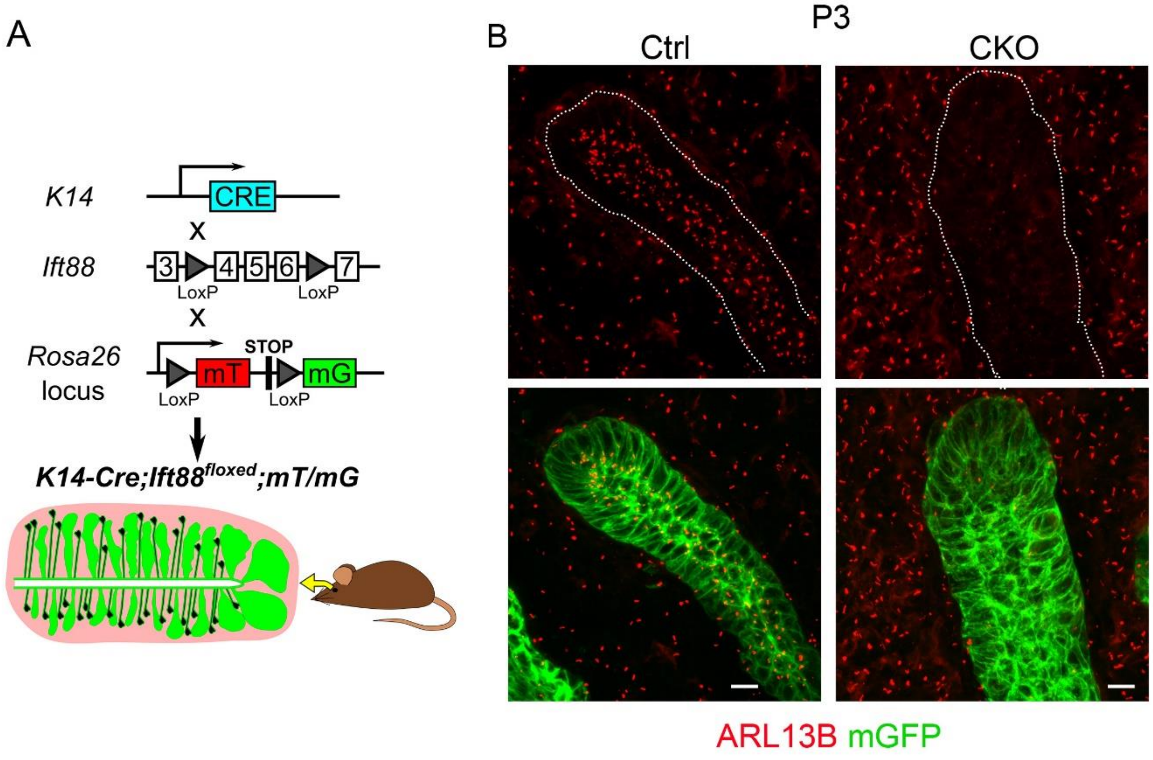


Figure S1

Bar= 10μm

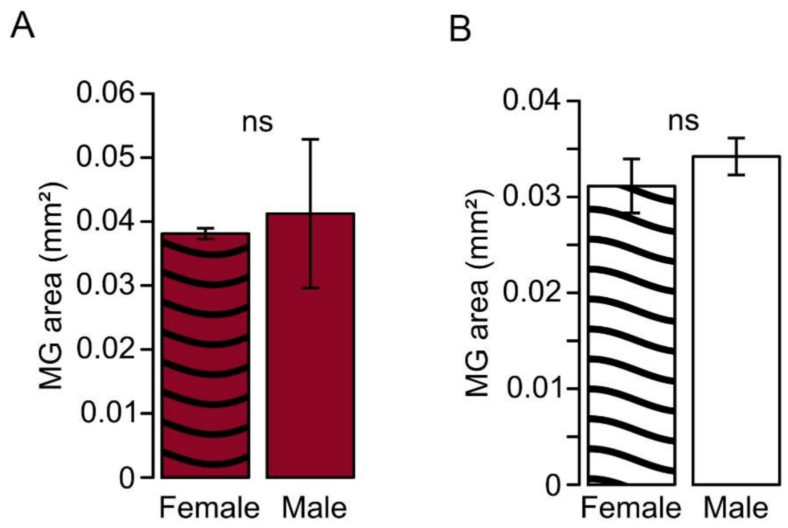

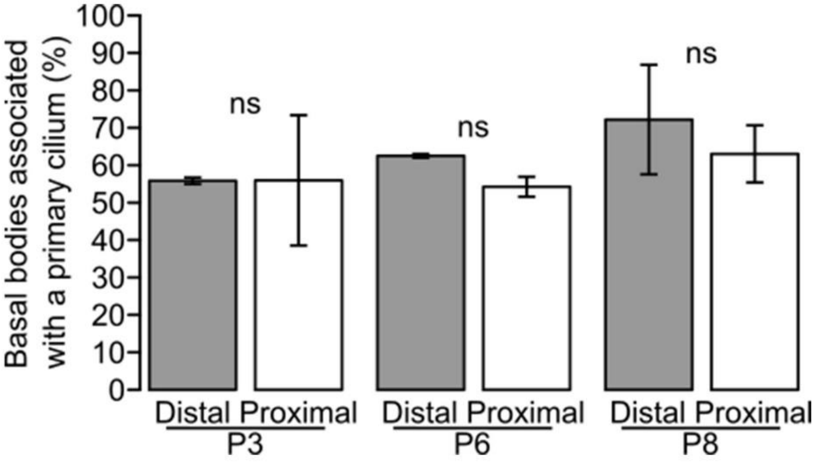
